# Bivalent, Copper-Mediated Amyloid-β Binding Disrupts the Prion Protein’s Neuroprotective Conformation

**DOI:** 10.64898/2026.11.16.767124

**Authors:** Amanda Smart, Francesca Pavlovici, Kevin Singewald, Cody Marshak, Claire Carollo, Sam Kaplan, Glenn L. Millhauser

**Author notes:** Corresponding Author: Correspondence should be addressed to Glenn L. Millhauser.

## Abstract

Emerging evidence shows that the cellular prion protein (PrP^C^) is a major receptor for Amyloid-β (Aβ) and that this aberrant interaction may influence the progression of Alzheimer’s disease (AD). However, the molecular basis of this interaction remains poorly understood. Both proteins strongly bind Cu^2+^, and we previously discovered that PrP^C^, Aβ30, and Cu^2+^ form a ternary complex at equimolar ratios. Here, using electron paramagnetic resonance (EPR) and nuclear magnetic resonance (NMR) spectroscopy with isotopically labeled proteins, we characterize the stoichiometry, stability, and molecular interactions of this ternary complex. We show that the complex forms preferentially at substoichiometric to equimolar Cu^2+^ concentrations yet remains stable at high Cu^2+^ ratios. The formation of the ternary complex is bivalent, requiring the previously identified Aβ-binding domain on PrP^C^ (residues 95-110) along with mutual Cu^2+^ coordination to the N-terminal octapeptide repeat domain in PrP^C^. Line broadening observed in Aβ30 NMR experiments identifies its negatively charged N-terminal residues and central hydrophobic core (residues 17-21) as the primary PrP^C^ binding sites. Through histidine mutation studies, we determine Aβ30 coordinates Cu^2+^ primarily through His14. Critically, the ternary complex formation displaces Cu^2+^ from a specific His residue on the C-terminal domain of PrP^C^, disrupting the neuroprotective *cis* interdomain interaction that otherwise regulates the toxic effector N-terminal domain. Using segmentally isotopically ^15^N-labeled PrP^C^, we show that while the *cis* interaction is not fully eliminated by interaction with Aβ, it becomes a disfavored conformational state. Together, these findings demonstrate how Cu^2+^ reinforces the PrP^C^–Aβ interaction, in turn destabilizing PrP^C^’s neuroprotective fold, offering a molecular mechanism for Aβ accumulation at—and potential trafficking across—the plasma membrane.

## Introduction

Alzheimer’s disease (AD) is a devastating neurodegenerative disorder characterized by progressive synaptic dysfunction, neuronal death, and decline in emotional processing and cognitive abilities.^1,2^ The accumulation of extracellular amyloid-β (Aβ) plaques and intracellular neurofibrillary tangles in the brain are characteristic hallmarks of AD.^3^ In recent years, the cellular prion protein (PrP^C^) has emerged as an important cellular receptor that mediates Aβ toxicity.^4–6^ Our work has shown that Cu^2+^ may be crucially involved in this interaction.^7,8^ The PrP^C^-Aβ interaction has been implicated in Aβ aggregation, copper dysregulation, and aberrant cellular signaling.^9–13^ However, the molecular mechanisms underlying PrP^C^-Aβ interactions and their contribution to these pathological processes remain poorly understood.

Aβ is a 38-42 amino acid intrinsically disordered, aggregation-prone extracellular peptide formed through sequential cleavage of the transmembrane amyloid precursor protein (APP).^14^ Figure 1A shows the Aβ sequence with highlighted histidine residues implicated in Cu^2+^ coordination and the central hydrophobic core (CHC, residues 17-21) which is critical for aggregation and therefore a primary target for therapeutic intervention.^15,16^ At physiological pH, Aβ coordinates Cu^2+^ through two histidine residues, Asp1 carbonyl, and the N-terminal amine.^17–19^ Previous studies highlight how Cu^2+^ dysregulation is implicated in AD pathogenesis, specifically finding that amyloid plaques retrieved from AD patients show a 200-400 fold enrichment of copper compared to normal brain tissue.^20–23^ Furthermore, monomeric and oligomeric Aβ can reduce bound Cu^2+^ to Cu^+^, producing toxic reactive oxygen species (ROS).^18,24^ The interactions of monomeric Aβ with other AD-related proteins, and consequent toxic pathways, remain poorly understood as the intrinsically disordered peptide aggregates above ∼0.09-0.5 µM, depending on the fragment.^25,26^

**Figure 1.**
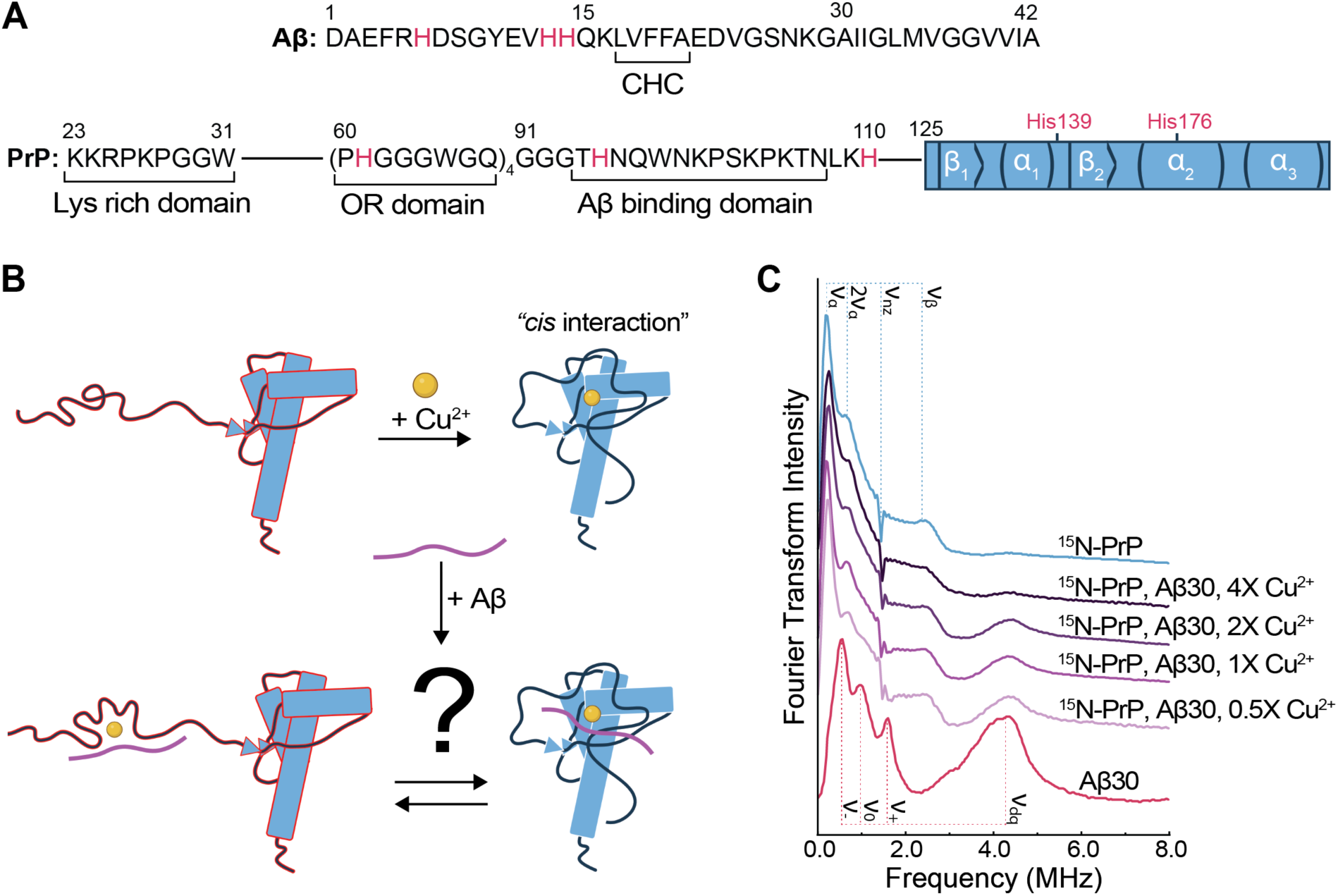
Cu^2+^ promotes formation of a stable ternary complex among Aβ30 and PrP^C^. **(A)** Linear sequences of Aβ (top) and PrP^C^ (bottom). Histidine residues involved in Cu^2+^ coordination are red. Both Aβ and the N-terminal domain of PrP^C^ (residues 23–125) are intrinsically disordered, whereas the PrP^C^ C-terminal domain (residues 126–230) is structured with secondary elements depicted in the blue box. The Aβ central hydrophobic core (CHC), residues 17-21 are indicated. The polybasic (PB), Cu^2+^-binding octapeptide repeat (OR) and Aβ binding domains are indicated in PrP^C^. **(B)** Schematic depicting structural changes in PrP^C^ (blue) upon interaction with Cu^2+^ (yellow circle) and Aβ (purple peptide). The open PrP^C^ conformation, associated with increased neurotoxicity, is indicated with a red outline. Figure created using BioRender. **(C)** Stacked ESEEM frequency spectra of 100 μM proteins coordinating to 1X Cu^2+^ at pH 7.4, unless otherwise stated. The ^15^N-His coordination frequencies (ν*_α_*, 2ν*_α_*, ν_nz_ and ν_β_) are labeled for ^15^N-PrP^C^ (blue), and the ^14^N-His coordination frequencies (ν_-_, ν_0_, ν_+_, and ν_dq_) are labeled for Aβ30 (red). Shown in purple (light to dark) are spectra of ^15^N-PrP^C^ with Aβ30 in the presence of 0.5–4 eq. of Cu^2+^. The ESEEM spectra have been normalized to their respective echo-intensities.

PrP^C^ is a cell surface glycoprotein expressed throughout the central nervous system.^27^ It plays important roles in synaptic function, cell signaling and metal ion homeostasis.^28^ Murine PrP^C^ is comprised of an intrinsically disordered N-terminal domain (residues 23-125) and a largely alpha-helical C-terminal domain (residues 126-231), illustrated in Figure 1A. The N-terminal domain contains three key regions of relevance here: the polybasic (PB) domain (residues 23-31), the octapeptide repeat (OR) domain (residues 59-90), and the Aβ-binding domain (residues 95-110).^4,29,30^ The Lys residues in the PB domain have also been shown to contribute to Aβ binding.^30^ The murine OR domain consists of four PHGGXWGQ repeats, where X is Gly in repeats one and four, or Ser in repeats two and three.^31^ At equimolar Cu^2+^, PrP^C^ coordinates Cu^2+^ through three histidine residues in the OR domain and one histidine on the surface of the C-terminal domain, with equilibrium favoring His176 over His139.^32,33^ This Cu^2+^ driven interdomain interaction, referred to as the *cis* interaction, is shown to be a neuroprotective conformation, as depicted in Figure 1B.^32,34^ When the *cis* interaction is disrupted, the N-terminal domain may extend away from the regulatory C-terminal domain, allowing the arginine-rich PB domain to penetrate the cell membrane leading to neurotoxic, transmembrane cationic currents.^34^ At saturation, PrP^C^ coordinates up to six Cu^2+^ ions: four at the OR domain (one per repeat), and two at non-OR N-terminal sites (His95 and His110).^35,36^ Interestingly, elevated Cu^2+^ concentrations promote PrP^C^ endocytosis, allowing for excess Cu^2+^ to be sequestered while PrP^C^ is recycled back to the cell surface.^28,37,38^

PrP^C^ is a receptor for oligomeric Aβ, an interaction that facilitates plaque formation^9^ and compromises synaptic function.^4^ Our lab showed that PrP^C^ also interacts with monomeric Aβ, as well as soluble Aβ30 (the peptide comprised of the first 30 residues) resulting in the peptide’s internalization through endocytosis.^7^ In our efforts to characterize the molecular details of the PrP^C^-Aβ interaction, we identified a ternary complex formed by PrP^C^, Aβ30, and Cu^2+^ at equimolar ratios and physiological pH.^8^ Formation and transport of this ternary complex would concentrate Aβ and Cu^2+^ in acidic vesicles, an environment capable of promoting Aβ and Aβ-Cu^2+^ aggregation.^39^ Moreover, Aβ could disrupt the Cu^2+^-driven *cis* interaction in PrP^C^ by displacing coordination from His176, thereby facilitating the previously observed membrane disruption and neurotoxicity, as shown schematically in Figure 1B.^34^ Thus, understanding the coordination and interaction between these key players could provide critical new infights into the underlying mechanisms of AD. Here, we characterize the stoichiometry, stability, and molecular details of the PrP^C^-Aβ30-Cu^2+^ ternary complex using complementary electron paramagnetic resonance (EPR) and nuclear magnetic resonance (NMR) spectroscopy experiments.

## Results and Discussion

### Copper drives formation of PrP^C^-Aβ ternary complex

Our recent work showed that PrP^C^, Aβ30 and Cu^2+^ form a ternary complex at equimolar ratios. Based on peak intensities in the electron spin echo-envelope modulation (ESEEM) EPR frequency domain, we found that Cu^2+^ coordinates to three histidine residues on PrP^C^ and a single histidine on Aβ30. This was surprising given the strong affinity of PrP^C^ for Cu^2+^, with K_d_ ranging from 0.3 nM-7.0 µM, taking up to 4 equivalents of Cu^2+^.^8,31,35,40^ By contrast, Aβ30 binds a single Cu^2+^ ion with an affinity of 16 µM at pH 7.4.^8,18,24^ To understand the constraints of this complex, we set out to first examine the Cu^2+^ binding limits of the complex. For this, we employed mixed isotope ESEEM EPR, a methodology recently developed by our lab.^8,41^ ESEEM probes the hyperfine coupling between the unpaired electron in Cu^2+^ and coupled nitrogen nuclei, and outputs characteristic frequency patterns that differ based on isotope.^8,42^ By using a mix of uniformly isotopically labeled ^15^N-PrP^C^ (I = 1/2) and natural abundance Aβ30 (I = 1), ESEEM can resolve each proteins coordination to Cu^2+^ as distinguished by the respective isotopes’ nuclear and associated spin transitions.^8,41^ Samples containing equimolar equivalents of ^15^N-PrP^C^ and Aβ30 were mixed with 0.5-4 eq. of Cu^2+^. The echo-intensity normalized ESEEM frequency domains are shown in Figure 1C. For reference, the individual proteins in complex with Cu^2+^ are also shown. Unless otherwise stated, all EPR experiments were performed with 100 µM protein at pH 7.4.

The frequency domain for ^15^N-PrP^C^ with 1 eq. Cu^2+^ shows 4 peaks, where ν*_α_*, 2ν*_α_* and ν_β_ are characteristic ^15^N-peaks and ν_nz_ corresponds to the ^15^N-nuclear Zeeman frequency.^42,43^ The Aβ30 with 1 eq. Cu^2+^ frequency domain shows the three nuclear quadrupole interactions (NQI) ν_-_, ν_0_, and ν_+_, and double quantum (DQ) ν_dq_ frequency peaks that correspond to ^14^N-His coordination.^44–46^ Both the presence of the 2ν*_α_* peak for ^15^N-His and the NQI: DQ peak intensity ratio for ^14^N-His can be used to asses when multiple histidine residues simultaneously coordinate to Cu^2+^.^8,47^ The ESEEM of ^15^N-PrP^C^, Aβ30 with 0.5, 1 or 2 eq. of Cu^2+^ show well-resolved ν*_α_*, 2ν*_α_*, ν_β_ and ν_dq_ peaks, recognizing that both ^15^N- and ^14^N-His residues are coordinating to Cu^2+^. The three spectra overlay well, suggesting that the complex is stable over a range of copper concentrations. Notably, the ternary complex formed to full occupancy even at substoichiometric Cu^2+^ levels despite PrP^C^ having a 32-fold higher binding affinity than Aβ30. The presence of the 2ν*_α_* peak and the low ν_dq_ peak intensity in the 0.5, 1 and 2 eq. Cu^2+^ ternary complex follows our previously reported ternary complex model and simulations that suggested a single Aβ30 His and multiple ^15^N-PrP^C^ His simultaneously coordinate to Cu^2+^.^8^ Although the ^15^N-His 2ν*_α_* peak is resolved in these three spectra, it can be distorted due to overlap with the low frequency ^14^N-NQI peaks. At 4 eq. of Cu^2+^ the signal looks very similar to that of ^15^N-PrP^C^ alone with 4 eq. of Cu^2+^, as seen in Figure S1 (spectra of ^15^N-PrP^C^ with 1-4 eq. Cu^2+^). However, there does still appear to be a very small ν_dq_ peak around 4 MHz, possibly indicating that Aβ30 is still able to coordinate even at high Cu^2+^ concentrations. Collectively, the Cu^2+^ titration suggests that PrP^C^, Aβ30, and Cu^2+^ form a stable ternary complex at 0.5-2 eq. Cu^2+^.

### PrP^C^ shifts Cu^2+^ coordination within Aβ30 upon ternary complex formation

Although ESEEM, as applied here, is an excellent tool for characterizing Cu^2+^-His coordination environments, it does not directly reveal interactions among the protein surfaces. To probe whether other regions on Aβ30 are binding to PrP^C^ in the ternary complex, we employed NMR spectroscopy. Our lab has previously characterized the PrP^C^-Cu^2+^ and PrP^C^-Aβ30 interactions using ^1^H-^15^N-Heteronuclear Single-Quantum Correlation (HSQC) NMR spectroscopy.^7,32,33^ To determine interacting domains and changes in conformation, we examined changes in the ^1^H-^15^N-HSQC peak intensities of uniformly labeled ^15^N-Aβ30. Peak intensity changes may arise from chemical/conformational exchange or, when a paramagnetic species such as Cu^2+^ is proximal to residues, paramagnetic relaxation enhancement (PRE).^48,49^

We first assigned ^15^N-Aβ30 HSQC peaks using previously obtained NOESY and TOCSY data, then compared assignments to published monomeric Aβ fragment spectra.^50,51^ We then characterized Aβ30 interactions with PrP^C^ and Cu^2+^ individually before examining the ternary complex. Figure 2A shows the HSQC spectrum of ^15^N-Aβ30 with PrP^C^ (black). Normalized peak intensity ratio (I/I_o_) bar charts of ^15^N-Aβ30 with PrP^C^ are shown in Figure 2B. The intensity ratios are calculated by dividing the HSQC-measured residue peak intensity of the experimental ^15^N-Aβ30 with PrP^C^ by the sample of ^15^N-Aβ30 alone and then scaled the peak maximum. Intensity ratios reveal that PrP^C^ binding results ∼50% signal loss across the first two-thirds of Aβ30, with residues 16-21 the most dramatically affected. The observed peak intensity reduction, along with line broadening, but without change to chemical shift indicates intermediate exchange. The first half of Aβ30 contains a cluster of negatively charged residues (D1, E3, D7, and E11) that could interact with the polybasic residues of PrP^C^, followed by the CHC (residues 17-21), which potentially interact with PrP^C^ hydrophobic segments. This finding is particularly significant given our previous observation that Aβ(1-16) and Aβ(16-30) did not elicit PrP^C^-mediated cellular uptake, yet Aβ30 was sufficient, further suggesting that not just charged or not just hydrophobic regions are enough for the interaction. These results agree with previous studies on PrP^C^ showing that both the highly charged polybasic domains and hydrophobic regions are necessary for interaction with Aβ.^4,29,30^ Notably, residues 17-21 comprise the CHC, which drives Aβ aggregation and is a major therapeutic target.^15,16^

**Figure 2.**
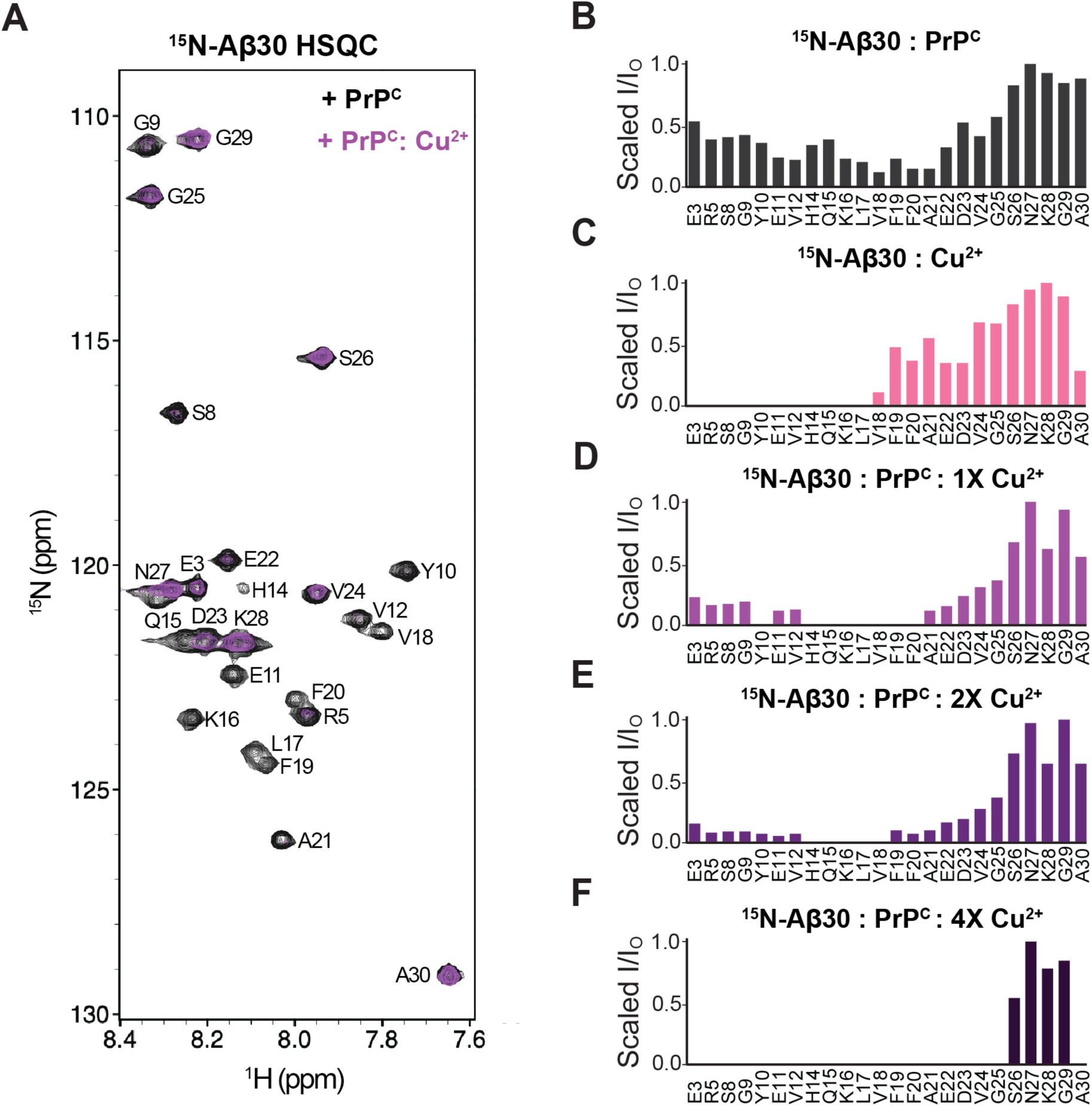
Aβ30 coordinates Cu^2+^ through central histidine residues in ternary complex. **(A)** Overlay of ^1^H–^15^N HSQC spectra of ^15^N-Aβ30 with PrP^C^ (black) and plus 1 eq. Cu^2+^ (purple). Normalized intensity ratio chart of ^15^N-Aβ30 in the presence of PrP^C^ **(B)**, Cu^2+^ **(C)**, PrP^C^ and 1 eq. Cu^2+^ **(D)**, PrP^C^ and 2 eq. Cu^2+^ **(E)**, and PrP^C^ and 4 eq. Cu^2+^ **(F)**, each compared to ^15^N-Aβ30. The intensity ratios are individually scaled and the assigned ^15^N-Aβ30 residues in the HSQC are labeled on the x-axis.

Next, we examined how ^15^N-Aβ30 is influence by Cu^2+^. The intensity ratios show dramatic signal loss from residues 1-17 due to Cu^2+^-induced PRE. This is consistent with the Aβ30-Cu^2+^ binding model, in which the N-terminal amine, Asp1 carbonyl, and two histidine residues (His6, His13, or His14) coordinate Cu^2+^. Finally, we added 1 eq. PrP^C^ and Cu^2+^ (1, 2, or 4 eq.) to ^15^N-Aβ30 to form the ternary complex. The single equivalent of Cu^2+^ in the ternary complex HSQC (purple) is overlayed on the ^15^N-Aβ30: PrP^C^ HSQC (black), in Figure 2A. The loss of several peaks indicates Cu^2+^ is interacting directly with Aβ in the PrP^C^-Aβ30 complex, consistent with the findings from the ESEEM experiments described above. However, forming the ternary complex shifts Cu^2+^ coordination toward the peptide center, as evidenced by signal loss in residues 14-20 and partial recovery of N-terminal peaks 3-12. This data points to His13 or His14 as the primary Cu^2+^ coordinating residue in the ternary complex. Consistent with ESEEM results, increasing Cu^2+^ from 1 to 2 eq. does not significantly alter the Cu^2+^ coordination environment around Aβ30. However, 4 eq. Cu^2+^ produces intense PREs across most of the peptide. Given that the ESEEM of this sample did not have an increase in ν_dq_, it would suggest that Aβ30 remains bound to PrP^C^ while additional Cu^2+^ ions coordinate to nearby PrP^C^ sites, leading to dramatic signal loss observed in the HSQC. However, this was not further tested. Collectively, these results demonstrate that the ternary complex forms across multiple Cu^2+^ concentrations and most likely involves His13 or His14 from Aβ30; this is further tested by mutagenesis experiments.

### Aβ30 Coordinates Cu^2+^ primarily through His14 in the ternary complex

To identify which Aβ30 histidine imidazole coordinates Cu^2+^ in the ternary complex and determine whether Aβ30-PrP^C^ binding is essential for complex formation, we employed site-directed mutagenesis. We generated single, double, and triple His to Ala mutations of Aβ30 at His6, His13, and His14, for a total of seven mutants. The His to Ala mutation was chosen to eliminate Cu^2+^ coordination while maintaining peptide solubility and preserving the intrinsically disordered state of Aβ30, as confirmed by CD spectroscopy (Figure S2). ESEEM analysis of Aβ30-Cu^2+^ coordination in the absence of PrP^C^ (Figure S2) revealed that single His mutations minimally affected Cu^2+^ binding. However, double mutations eliminating His14 (Aβ30 H6A/H14A and H13A/H14A) dramatically reduced Cu^2+^ binding and altered coordination geometry, establishing His14 as essential for Aβ30-Cu^2+^ coordination.

Having characterized how each mutation affects Aβ30-Cu^2+^ coordination, we next examined ternary complex formation. Figure 3A shows ESEEM of ternary complexes with each mutant, excluding the triple mutant (Aβ30 H6A/H13A/H14A, hereafter referred to as Aβ30 ΔHis), which fully eliminates ^14^N-His coordination. Surprisingly, most mutant ternary complexes showed similar ν*_α_* and ν_dq_ peak intensities, suggesting that any His within Aβ30 may participate in Cu^2+^-coordination. However, the 2ν*_α_* and NQI overlapping domains showed subtle changes among samples, possibly indicating minor shifts in coordination geometry while maintaining the 3:1 ^15^N-PrP^C^-His:^14^N-Aβ30-His ratio. Only the H6A/H14A double mutant showed decreased ν_dq_ intensity, indicating His13 is less preferred in the ternary complex. However, the consistent ESEEM signals across mutants suggest that the ternary complex can form with any Aβ30 histidine, but with preference for His6 or His14.

**Figure 3.**
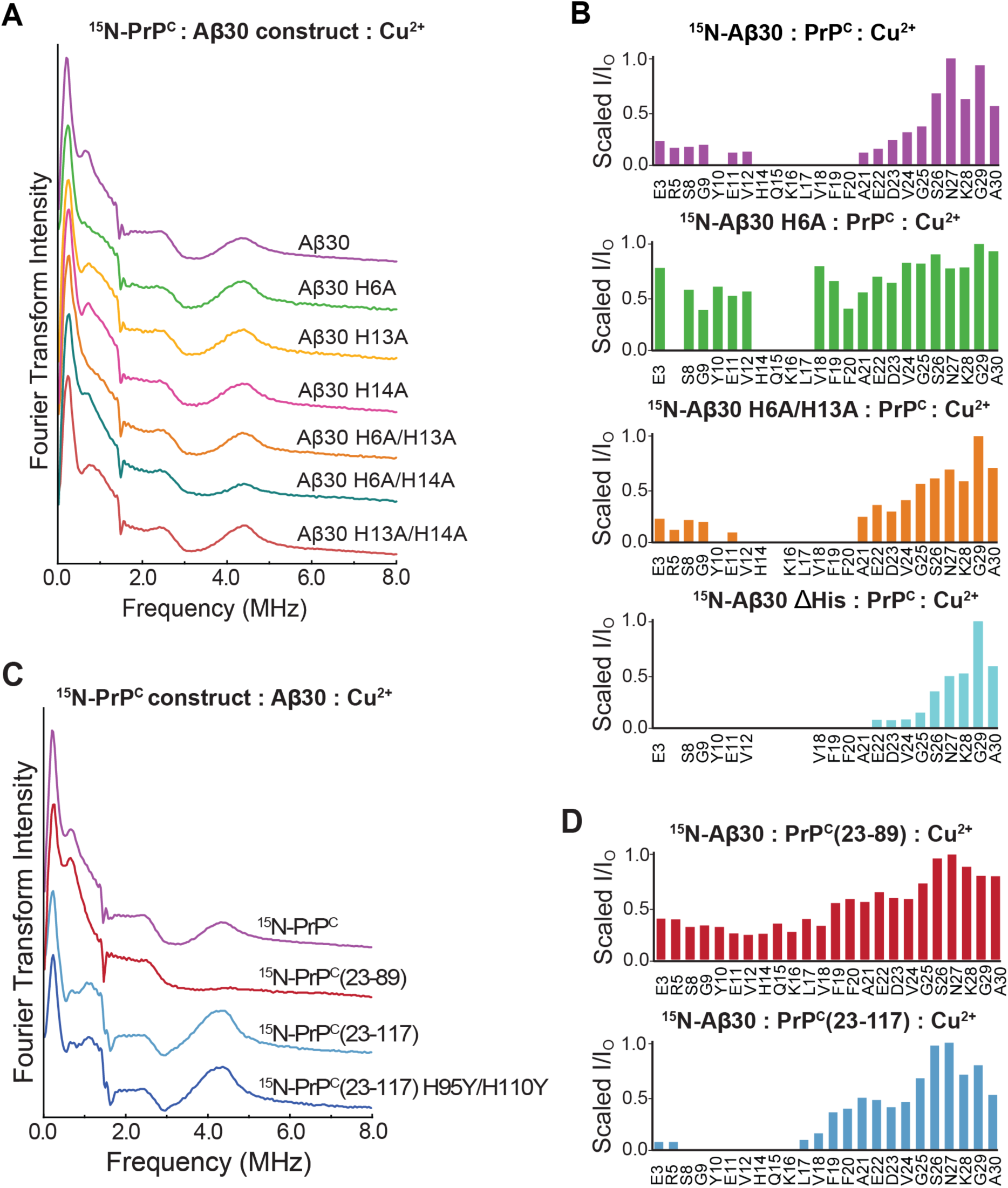
Aβ30 primarily coordinates Cu^2+^ through His14 in ternary complex and requires the PrP^C^-Aβ binding. **(A)** Stacked ESEEM frequency domain spectra of ^15^N-PrP^C^ and Cu^2+^ with Aβ30 (purple), Aβ30 H6A (green), Aβ30 H13A (yellow), Aβ30 H14A (pink), Aβ30 H6A/H13A (orange), Aβ30 H6A/H14A (turquoise), or Aβ30 H13A/H14A (dark orange). **(B)** Normalized intensity ratios from HSQC of ¹⁵N-Aβ30 (purple), ¹⁵N-Aβ30 H6A (green), ¹⁵N-Aβ30 H6A/H13A (orange), or ¹⁵N-Aβ30 ΔHis (cyan) in the presence of PrP^C^ and Cu^2+^. The measurable residue peaks for each ¹⁵N-Aβ30 species are shown on the x-axis and aligned with the wildtype sequence. **(C)** Stacked ESEEM frequency spectra of Aβ30 and Cu^2+^ with ^15^N-PrP^C^ (purple), ^15^N-PrP^C^(23-89) (red), ^15^N-PrP^C^(23-117) (blue), or ^15^N-PrP^C^(23-117) H95Y/H110Y (dark blue). **(D)** HSQC normalized intensity ratios of ¹⁵N-Aβ30 and Cu^2+^ with ^15^N-PrP^C^(23-89) (red) or ^15^N-PrP^C^(23-117) (blue).

To map coordination changes at the Aβ30 interface, we next performed HSQC NMR on select His to Ala mutants. Since ESEEM showed only subtle differences in ternary complex Cu^2+^ coordination among mutants, we focused on ^15^N-Aβ30 H6A, ^15^N-Aβ30 H6A/H13, and ^15^N-Aβ30 ΔHis. Intensity ratios comparing these mutants to the WT ternary complex are shown in Figure 3B, with additional NMR double mutant data in Figure S3. As Aβ30 is intrinsically disordered, point mutations cause chemical shift perturbations in nearby residues, reducing peak identification in some spectra. By contrast with the ESEEM experiments, HSQC revealed distinct changes in the Aβ30-Cu^2+^ interaction. The Aβ30 H6A ternary complex showed complete N-terminal peak recovery with PREs localized to residues near His13 and His14. Because ESEEM suggested His13 is unfavorable, we examined the Aβ30 H6A/H13A double mutant to determine whether His14 alone mediates ternary complex formation like WT. This spectrum resembled the WT ternary complex, with intense Cu^2+^-induced PREs at residues 12-20 causing complete signal loss, thereby confirming His14 as the primary Cu^2+^-coordinating residue. Lastly, we examined the ^15^N-Aβ30 ΔHis mutant in the ternary complex. Remarkably, intense PREs affected residues 3-20 despite Aβ30 ΔHis mutant fully lacking His residues. This supports that Aβ30 binds PrP^C^ proximal to the Cu^2+^ coordination site, allowing Cu^2+^-induced PREs to affect Aβ30 residues even without direct Cu^2+^ coordination. The altered intensity ratios also likely reflect changes in PrP^C^-Aβ30 interaction, due to the change in peptide hydrophobicity as reflected by the change in intensity ratios for ^15^N-Aβ30 ΔHis with PrP^C^ (Figure S3). It is also possible that this mutant coordinates to Cu^2+^ through its N-terminal amine, although typically unfavored at a pH of 7.4.^18^ This was not examined further.

We next investigated whether the Aβ-binding domain in PrP^C^ (residues 95-110) participates in the formation of the ternary complex. We expressed two N-terminal PrP^C^ constructs: PrP^C^(23-89) retaining the OR domain, and PrP^C^(23-117) extended to include residues 95-110.^4,33,52^ Both constructs bind Cu^2+^ normally as shown by EPR (Figure S4). The ESEEM spectrum of ^15^N-PrP^C^(23-89), Aβ30, and Cu^2+^ in Figure 3C shows no ν_dq_ peak, indicating Aβ30 cannot coordinate Cu^2+^. Consistent with this, the HSQC of ^15^N-Aβ30, PrP^C^(23-89), and Cu^2+^ shows only minor intensity changes across Aβ30. By contrast, ESEEM of ^15^N-PrP^C^(23-117), Aβ30, and Cu^2+^ shows a ν_dq_ peak, indicating the Aβ-binding domain biases towards ternary complex formation. Notably, the ν_dq_ peak intensity is higher than in the full-length ternary complex, suggesting Aβ30 now contributes two histidine residues to Cu^2+^ coordination.

To determine whether Aβ30 coordinates Cu^2+^ at the OR domain on PrP^C^ or at the non-OR sites (His95/His110), we generated PrP^C^(23-117) H95Y/H110Y double mutant. ESEEM of the ^15^N-PrP^C^(23-117) H95Y/H110Y mutant remained identical to ^15^N-PrP^C^(23-117), confirming Aβ30 coordinates Cu^2+^ within the OR domain. The HSQC of ^15^N-Aβ30 with PrP^C^(23-117) and Cu^2+^ resembles the full-length ternary complex. These results demonstrate that the canonical ternary complex requires that Aβ interact with PrP^C^ through both mutual Cu^2+^ coordination and protein-protein contacts involving PrP^C^ residues 95-110.

### Formation of the ternary complex disrupts the neuroprotective cis interaction in PrP^C^

As noted in the introduction, Cu^2+^ suppresses inherent PrP^C^ neurotoxicity by simultaneously coordinating to His residues in the OR domain and His176 in the regulatory C-terminal domain. Given the ternary complex with Aβ requires one of the Aβ His residues, we questioned whether this interaction might displace His176. Previously published studies of ^15^N-PrP^C^ with 1 eq. Cu^2+^ identified three patches on the C-terminal domain with dramatically reduced peak intensities due to Cu^2+^-induced PREs.^32^ Patches 1 and 2 are centered on His139 and His176, respectively, and indicate *cis* interaction formation. Patch 3 overlays the acidic surface on the C-terminal domain and further contributes to the *cis* interaction through electrostatic attraction with the OR-Cu^2+^ center. To test whether Aβ30 affects PrP^C^-Cu^2+^ interactions, we first examined ^15^N-PrP^C^ with varying Cu^2+^ concentrations. Figure 4A shows the normalized peak intensity ratios of ^15^N-PrP^C^ with 1, 2, or 4 eq. Cu^2+^. As the Cu^2+^ concentration increases, we observed systematic decreases in peak intensities across each of the three C-terminal patches.

**Figure 4.**
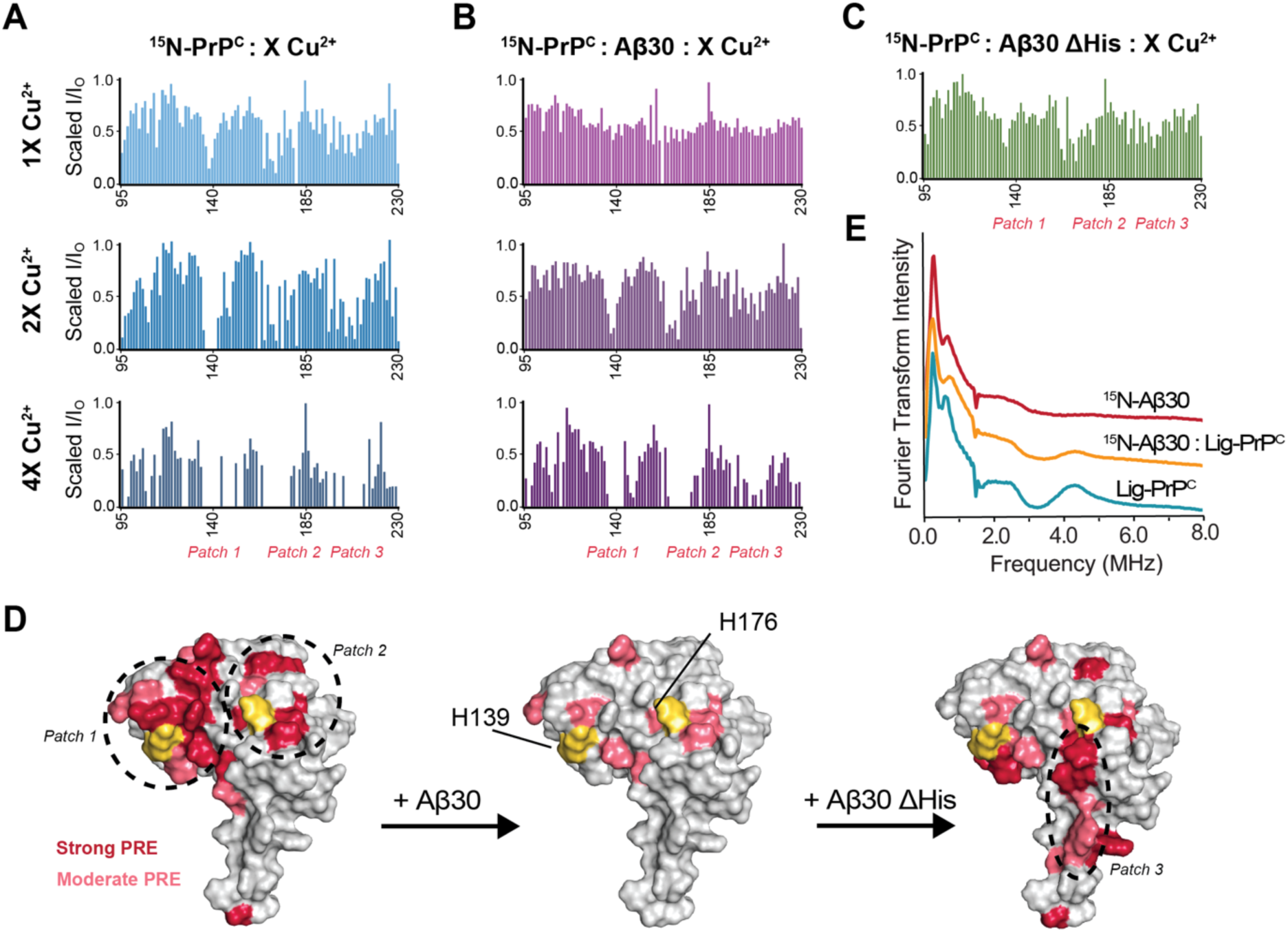
Aβ30 localizes Cu^2+^ at PrP^C^ N-terminal domain disrupting the *cis* interaction. **(A)** Normalized intensity ratios of ^15^N-PrP^C^ in the presence of one (light blue), two (blue), or four (dark blue) equivalents of Cu^2+^, compared to ¹⁵N-PrP^C^. **(B)** Normalized intensity ratios of ^15^N-PrP^C^ in the presence of Aβ30 and one (light purple), two (purple), or four (dark purple) equivalents of Cu^2+^, compared to ^15^N-PrP^C^. **(C)** Normalized intensity ratios of ^15^N-PrP^C^ in the presence of Aβ30 ΔHis and one equivalent of Cu^2+^, compared to ^15^N-PrP^C^. **(D)** Surface representation of the PrP^C^ C-terminal domain (PDB: 1XYX) showing residues significantly affected by Cu²⁺-induced PREs with 1 eq. Cu^2+^ (left), and the addition of Aβ30 (middle) or Aβ30 ΔHis (right). Residues strongly affected (>3 MAD) are shown in red; moderately affected residues (2–3 MAD) are shown in pink; and C-terminal His139 and His176 are highlighted in yellow. **(E)** Stacked ESEEM frequency spectra of Cu^2+^ with ^15^N-Aβ30 (red), ^15^N-Aβ30 and Lig-PrP^C^ (pink) or Lig-PrP^C^ (teal).

Next, we repeated the experiment in Figure 4A but with the addition of one equivalent of Aβ30. The results, displayed in Figure 4B, show a significant reduction of the PRE effect from Cu^2+^. Specifically, the experiment with 1.0 eq. of Cu^2+^ exhibits few strongly affected residues, suggesting that Aβ30 displaces Cu^2+^ from the C-terminal domain. Experiments at 2.0 and 4.0 eq. of Cu^2+^ do show the three identifiable patches of reduced peak intensities, indicating a shift in the conformational equilibrium favoring the *cis* interaction. To distinguish whether *cis* interaction disruption results from Aβ30-PrP^C^ binding or Aβ30-Cu^2+^ coordination, we examined complexes formed with Aβ30 ΔHis (Figure 4C). The lack of any discernable, systematic influence on the PrP^C^ I/I_o_ values suggests that Aβ30 His residues are essential for promoting the Cu^2+^-mediated interaction and disruption to the *cis* interaction.

To visualize and systematically characterize the impact of Cu^2+^ displacement from the C-terminal domain of PrP^C^, Figure 4D shows peaks statistically affected by Cu^2+^-induced PREs mapped onto the surface representation of PrP^C^ (PDB: 1XYX) at one equivalent.^53^ The two C-terminal histidine residues, His139 and His176, are marked in yellow and are surrounded by residues significantly affected by Cu^2+^-induced PREs (>3 MAD, median absolute deviation), colored red, and residues moderately affected (2-3 MAD), colored pink. The drastic reduction in significantly affected residues by Cu^2+^ induced PREs at the C-terminal domain with the addition of Aβ30 clearly disfavors the PrP^C^ *cis* conformation. However, when we substitute Aβ30 with Aβ30 ΔHis, the *cis* interaction is partially recovered, though not to the level observed in the absence of Aβ30. Instead, it appears that the influence of Cu^2+^ moves closer to Patch 3, similar to patterns we have seen when both His139 and His176 were mutated to tyrosine and unable to bind Cu^2+^. ^32^Aβ30 therefore causes a change in how PrP^C^ coordinates Cu^2+^ and the formation of the *cis* interaction, even when it cannot directly coordinate Cu^2+^.

To definitively test whether the *cis* interaction persists in the ternary complex, we employed segmentally isotopically labeled PrP^C^ (Lig-PrP^C^), in which residues 23-117 are uniformly ^15^N-labeled while residues 118-230 retain natural-abundance ^14^N. This construct enables independent detection of N-terminal (^15^N) and C-terminal (^14^N) Cu^2+^ coordination by ESEEM spectroscopy.^52^ Figure 4E shows the ESEEM spectrum of Lig-PrP^C^ with 1 eq. Cu^2+^. Frequency peaks corresponding to ^15^N-OR histidine residues and ^14^N-C-terminal histidine residues coordinating Cu^2+^ are both present, demonstrating simultaneous coordination to the metal ion center. Addition of the ^15^N-Aβ30 results in a loss of signal intensity at ∼4.0 MHz, indicating a shift away from Cu^2+^ coordination from a PrP^C^ C-terminal His. The 4.0 MHz signal is not fully eliminated, thereby demonstrating that Aβ30 disrupts but does not eliminate the Cu^2+^-driving *cis* interaction. Collectively, these results demonstrate that the neuroprotective *cis* interaction can still form in the presence of Aβ30, but it is not the predominant state. Instead, ternary complex formation at the OR domain becomes the favored conformation, effectively liberating the N-terminal domain on PrP^C^. As such, monomeric Aβ could drive a deleterious pathway by potentiating the toxic effector PrP^C^ N-terminal domain.

## Discussion

By identifying the participating Aβ His residues, as well as protein-protein contacts separate from mutual ion coordination, this study extends our previous work that demonstrated a ternary complex among PrP^C^, Aβ30 and Cu^2+^. As established elsewhere, the polybasic residues of PrP^C^ in segments 23-31 and 95-110, including hydrophobic residues within 95-110, are contact points for Aβ binding.^4,29,30^ NMR experiments here show that residues 1-21 of Aβ30 participate in PrP^C^ binding. These include four negatively charged residues (D1, E3, D7, E11) and the central hydrophobic core (CHC, residues 17-21), which would engage the Aβ-binding domain of PrP^C^. Defining the specific contact points involved in the ternary complex is more challenging because Cu^2+^ induced PRE-based line broadening is not residue specific. Nonetheless, we showed that the Aβ-binding domain on PrP^C^ (residues 95-110) is required for ternary complex formation, demonstrating that protein-protein interaction is essential, contributing an additional binding site beyond the metal-bridged complex. Our findings support the hypothesized ternary complex model, in Figure 5.

**Figure 5.**
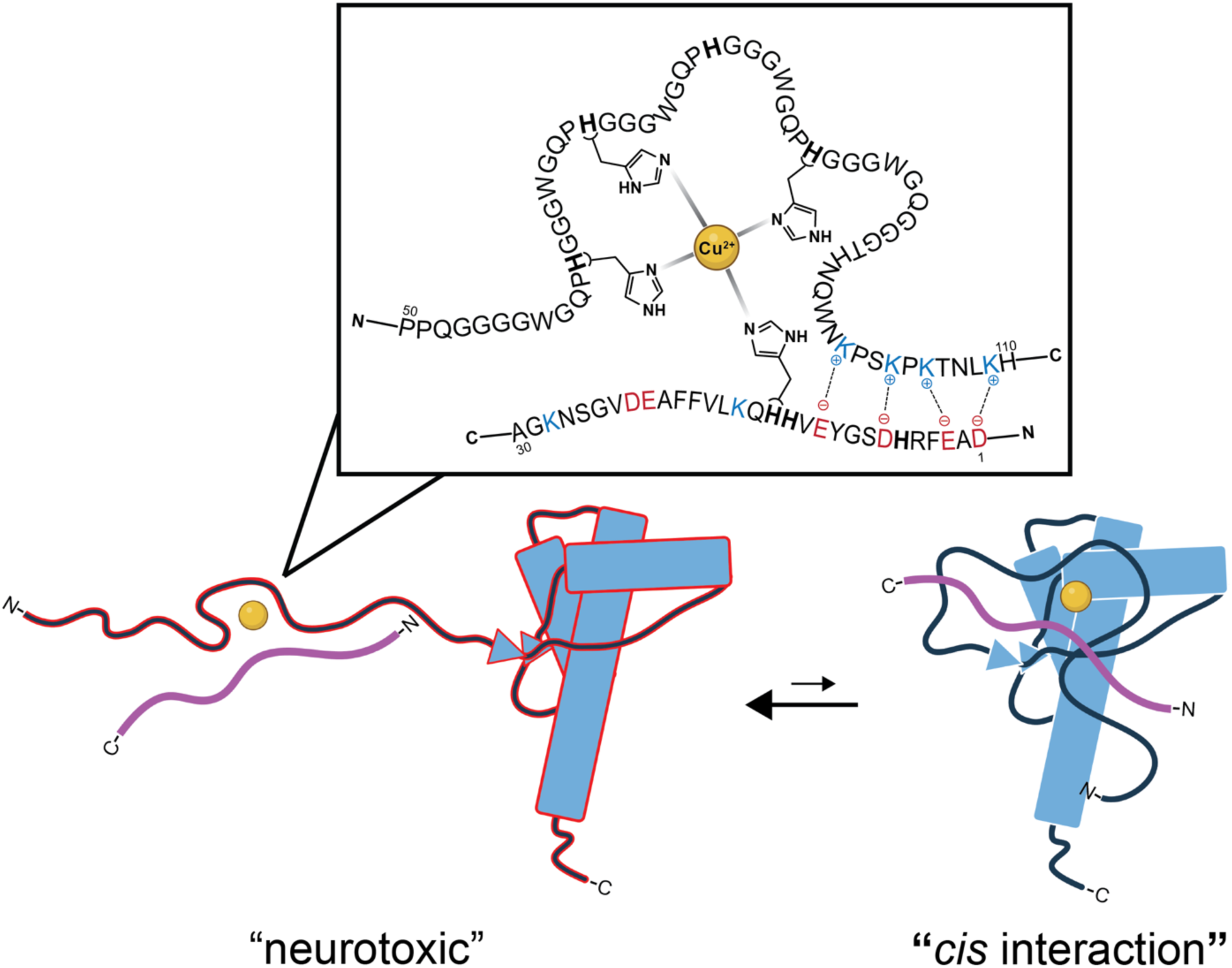
Model of ternary complex states. The schematic structure of PrP^C^ is shown in blue with the unstructured, toxic effector N-terminal domain unregulated (left) or regulated (right) by the structured C-terminal domain via Cu^2+^. The unstructured Aβ30 protein is shown in purple and interreacting with both Cu^2+^ and PrP^C^. The findings here support the model on the left as dominant. The bivalent interaction involving histidine-Cu^2+^ coordination and interacting charged residues on PrP^C^(50-110) and Aβ30 are shown in the insert.

In characterizing the ternary complex metal coordination, we showed that Aβ30 primarily engages Cu^2+^ through His14, with His6 and His13 serving as secondary contributors. The complex involves the PrP^C^ OR and C-terminal histidine residues, but not His95 or His110, which flank the Aβ-binding domain. The ternary complex forms within a range of Cu^2+^ concentrations, stable from 0.5-4 eq. As demonstrated with segmentally isotopically labeled PrP^C^, the *cis* interaction is not eliminated but becomes a disfavored conformational state in the presence of Aβ30. Even when Aβ30 cannot directly bind Cu^2+^, as in the Aβ30 ΔHis mutant, the *cis* interaction is partially impacted. This suggests that Aβ, in the formation of the ternary complex, alters how PrP^C^ interacts with copper, potentially altering its functionality at the cell surface, perhaps by influencing normal C-driven endocytosis.

Overall, these findings reveal a copper-facilitated mechanism by which PrP^C^ and Aβ bind, providing a molecular framework for understanding how this interaction drives monomeric and oligomeric Aβ toxicity. While our study used monomeric Aβ30, we previously showed that Aβ30 recapitulates PrP^C^-dependent uptake of full-length Aβ40 in cells,^7^ suggesting these findings may extend to Aβ42, as well as oligomeric Aβ, a primary toxic species that initiates AD. Our results are consistent with a Cu^2+^-mediated uptake pathway for Aβ, as established for PrP^C^ endocytosis,^37,38^ in which PrP^C^ facilitates endocytosis of the ternary complex, concentrating Aβ and Cu^2+^ in acidic lysosomes. This acidic environment may promote rapid Aβ aggregation and allow these aggregates to sequester Cu^2+^, potentially explaining the copper enrichment observed in amyloid plaques.^20,22,23^ Disrupting this copper-bridged complex could offer a strategy for inhibiting Aβ cellular uptake, thereby reducing Aβ aggregation. Future work must explore whether targeting this ternary complex can effectively reduce disease progression, thus providing a novel and certainly unexplored therapeutic strategy for Alzheimer’s disease.

## Experimental procedures

### Protein Expression and Purification

The prion protein was expressed following a previously established protocol.^33^ Briefly, a *M. musculus* PrP(23-230) construct cloned into the pJ414 vector (DNA 2.0) was transformed into *E. coli* BL21(DE3) (Invitrogen) for expression. Cultures were grown in LuriaBroth (Research Products International) or in M9 minimal media supplemented with ¹⁵N ammonium chloride (1 g/L, Cambridge Isotopes) to produce uniformly ¹⁵N-labeled protein. Cultures were maintained at 37 °C until reaching an OD₆₀₀ of 1.0, at which point expression was induced by addition of 1 mM IPTG and continued overnight at 25 °C. Inclusion bodies were solubilized at room temperature in a denaturing buffer containing 8 M guanidinium chloride (GndHCl), 100 mM Tris, and 100 mM sodium acetate at pH 8. The clarified soluble fraction was subjected to Ni²⁺-IMAC purification, with protein eluted using 5 M GndHCl, 100 mM Tris, and 100 mM sodium acetate at pH 4.5. Pooled fractions were brought to pH 8 using 6 M KOH and held at 4 °C for two days to permit oxidative formation of the native disulfide bond. Protein was subsequently desalted into 50 mM potassium acetate buffer and further resolved by reverse-phase liquid chromatography on a C8 column. The final purified PrP^C^ was lyophilized and stored at −70 °C.

Aβ30 was cloned into a pET-28b(+) (Novagen) vector encoding an N-terminal His-SUMO tag (provided by Carrie Partch, UCSC) via Gibson Assembly and expressed as previously described.^8^ Briefly, cells were cultured in Luria broth (Research Products International) or M9 minimal media supplemented with ¹⁵N ammonium chloride (1 g/L, Cambridge Isotopes) for uniformly ¹⁵N-labeled protein at 37 °C with shaking until OD₆₀₀ reached 0.6–0.8, then induced with 1 mM IPTG and grown overnight at 18 °C. Cultures were harvested, centrifuged and the cell pellets were resuspended in lysis buffer consisting of 50 mM Tris, 300 mM NaCl, 1 mM β-mercaptoethanol (βME), and Pierce Protease Inhibitor Tablets (Thermo Fisher Scientific) at pH 7.5 and sonicated. The soluble fraction was loaded onto Ni²⁺-IMAC resin at 4 °C and was washed sequentially with 60 mL lysis buffer and 30 mL cleavage buffer (50 mM Tris, 50 mM NaCl, 30 mM imidazole, 1 mM βME, pH 7.5). On-column cleavage of the His-SUMO tag was carried out by overnight incubation at 4 °C with 3 mM recombinantly expressed His-ULP1 protease. Aβ30 was recovered by elution buffer (50 mM Tris, 50 mM imidazole, 1 mM βME, pH 7.5) and further purified by reverse-phase HPLC on an Agilent PLRP-S column under basic conditions. Purified Aβ30 was lyophilized and stored at −70 °C.

### Pulsed Electron Paramagnetic Resonance

Lyophilized PrP^C^ was reconstituted in water, while Aβ30 was first dissolved in a small volume of 20 mM KOH before dilution with water. Protein concentrations were determined spectrophotometrically from absorbance at 280 nm using molar extinction coefficients of 63,495.0 M⁻¹·cm⁻¹ and 1,490.0 M⁻¹·cm⁻¹ for PrP^C^ and Aβ30, respectively Samples were prepared at 100 µM for each protein with ranging concentrations of Cu^2+^ in 25 mM HEPES containing 20% glycerol as a cryoprotectant, and pH was adjusted to 7.4 using 100 mM KOH. Samples were equilibrated at room temperature for a minimum of 15 minutes prior to freezing in 4 mm quartz X-band tubes (Wilmad-LabGlass).

ESEEM experiments were acquired on a Bruker E580 X-band (∼9.7 GHz) pulsed EPR spectrometer equipped with a MD4 resonator. All pulsed measurements were performed at the maximum of the Cu^2+^ absorption envelope at 3316 G and at a temperature of 18 K to maximize signal-to-noise for Cu^2+^ detection. The three-pulse ESEEM sequence (π/2–τ–π/2–T–π/2–τ–echo) employed four-step phase cycling. The interpulse delay τ was set to the hydrogen blind spot condition, calculated to be 210 ns at the applied magnetic field. T was incremented from an initial value of 12 ns in steps of 16 ns over 602 points, with a π/2 pulse width of 8 ns. Raw ESEEM time-domain data were intensity-normalized, background-corrected by exponential decay subtraction, zero-filled to 2048 points, and Fourier transformed using Bruker XEPR software.

### NMR Spectroscopy

All proteins were dissolved as described in preparing the EPR samples. The NMR samples were prepared at a final protein concentration of 100 µM in 25 mM MES (2-(N-morpholino)ethanesulfonic acid, Sigma) buffer adjusted to pH 6.6 with 1 M HCl and supplemented with 10% D₂O. Both ^1^H and ^1^H-^15^N HSQC spectra were collected on a Bruker 800 MHz spectrometer at the UCSC NMR Facility (Santa Cruz, CA). Spectral data were processed and analyzed using NMRPipe and POKY.

### Associated Content

The Supporting information is available.

## Author Contributions

**Amanda Smart**: Conceptualization (Lead), Methodology (Lead), Validation (Lead), Formal Analysis (Lead), Investigation (Lead), Resources (Supporting), Data Curation (Lead), Writing – Original draft (Lead), Writing – Review & Editing (Equal), Visualization (Lead), and Project Administration (Equal). **Glenn L. Millhauser**: Conceptualization (Supporting), Validation (Supporting), Resources (Lead), Writing – Original draft (Equal), Writing – Review & Editing (Equal), Supervision (Lead), Project Administration (Equal), and Funding acquisition (Lead).

## Acknowledgement

This research was supported by the National Institutes of Health grants R35GM131781 (G.L.M), the Cinvestav UC Alianza MX (G.L.M.), and the Initiative for Maximizing Student Development Program (1T32GM135742-02, A.S.). The pulsed EPR instrument was funded through NIH grant S10OD024980 and the 800 MHz NMR spectrometer was funded through the S10OD018455. We thank Dr. Jack Lee, Dr. Eefei Chen and, Dr. Victor E López-Guerrero for helpful conversation.

## Abbreviations

Aβ30: amyloid-β residues 1-30
AD: Alzheimer’s disease
CDC: central hydrophobic core
CW: continuous wave
DQ: double quantum
EPR: electron paramagnetic resonance
ESEEM: electron spin echo envelope modulation
HSQC: heteronuclear single quantum coherence
IDP: intrinsically disordered peptide
NMR: nuclear magnetic resonance
NQI: nuclear quadrupole interaction
OR: octapeptide repeat
PRE: paramagnetic relaxation enhancement
PrP^C^: cellular isoform of prion protein residues 23-230

## REFERENCES

1. Selkoe, D.J. (2024). The advent of Alzheimer treatments will change the trajectory of human aging. Nat Aging 4, 453–463. 10.1038/s43587-024-00611-5.

2. Pradeepkiran, J.A., Baig, J., Islam, M.A., Kshirsagar, S., and Reddy, P.H. (2025). Amyloid-β and Phosphorylated Tau are the Key Biomarkers and Predictors of Alzheimer’s Disease. Aging and disease 16, 658. 10.14336/AD.2024.0286.

3. Jack, C.R., Andrews, J.S., Beach, T.G., Buracchio, T., Dunn, B., Graf, A., Hansson, O., Ho, C., Jagust, W., McDade, E., et al. (2024). Revised criteria for diagnosis and staging of Alzheimer’s disease: Alzheimer’s Association Workgroup. Alzheimer’s & Dementia 20, 5143–5169. 10.1002/alz.13859.

4. Laurén, J., Gimbel, D.A., Nygaard, H.B., Gilbert, J.W., and Strittmatter, S.M. (2009). Cellular prion protein mediates impairment of synaptic plasticity by amyloid-β oligomers. Nature 457, 1128–1132. 10.1038/nature07761.

5. Smith, L.M., Kostylev, M.A., Lee, S., and Strittmatter, S.M. (2019). Systematic and standardized comparison of reported amyloid-β receptors for sufficiency, affinity, and Alzheimer’s disease relevance. Journal of Biological Chemistry 294, 6042–6053. 10.1074/jbc.RA118.006252.

6. Legname, G., and Scialò, C. (2020). On the role of the cellular prion protein in the uptake and signaling of pathological aggregates in neurodegenerative diseases. Prion 14, 257–270. 10.1080/19336896.2020.1854034.

7. Foley, A.R., Roseman, G.P., Chan, K., Smart, A., Finn, T.S., Yang, K., Lokey, R.S., Millhauser, G.L., and Raskatov, J.A. (2020). Evidence for aggregation-independent, PrP ^C^ - mediated Aβ cellular internalization. Proc Natl Acad Sci USA 117, 28625–28631. 10.1073/pnas.2009238117.

8. Smart, A., Singewald, K., Hasanbasri, Z., Britt, R.D., and Millhauser, G.L. (2025). Identifying the copper coordination environment between interacting neurodegenerative proteins: A new approach using pulsed EPR with 14N/15N isotopic labeling. Journal of Biological Chemistry 301, 108311. 10.1016/j.jbc.2025.108311.

9. Schwarze-Eicker, K., Keyvani, K., Görtz, N., Westaway, D., Sachser, N., and Paulus, W. (2005). Prion protein (PrPc) promotes β-amyloid plaque formation. Neurobiology of Aging 26, 1177–1182. 10.1016/j.neurobiolaging.2004.10.004.

10. You, H., Tsutsui, S., Hameed, S., Kannanayakal, T.J., Chen, L., Xia, P., Engbers, J.D.T., Lipton, S.A., Stys, P.K., and Zamponi, G.W. (2012). Aβ neurotoxicity depends on interactions between copper ions, prion protein, and N-methyl-D-aspartate receptors. Proc. Natl. Acad. Sci. U.S.A. 109, 1737–1742. 10.1073/pnas.1110789109.

11. Posadas, Y., Parra-Ojeda, L., Perez-Cruz, C., and Quintanar, L. (2021). Amyloid β Perturbs Cu(II) Binding to the Prion Protein in a Site-Specific Manner: Insights into Its Potential Neurotoxic Mechanisms. Inorg. Chem. 60, 8958–8972. 10.1021/acs.inorgchem.1c00846.

12. Haas, L.T., Kostylev, M.A., and Strittmatter, S.M. (2014). Therapeutic Molecules and Endogenous Ligands Regulate the Interaction between Brain Cellular Prion Protein (PrPC) and Metabotropic Glutamate Receptor 5 (mGluR5). Journal of Biological Chemistry 289, 28460–28477. 10.1074/jbc.M114.584342.

13. Larson, M., Sherman, M.A., Amar, F., Nuvolone, M., Schneider, J.A., Bennett, D.A., Aguzzi, A., and Lesné, S.E. (2012). The Complex PrP ^c^-Fyn Couples Human Oligomeric Aβ with Pathological Tau Changes in Alzheimer’s Disease. J. Neurosci. 32, 16857–16871. 10.1523/JNEUROSCI.1858-12.2012.

14. O’Brien, R.J., and Wong, P.C. (2011). Amyloid Precursor Protein Processing and Alzheimer’s Disease. Annu. Rev. Neurosci. 34, 185–204. 10.1146/annurev-neuro-061010-113613.

15. De Groot, N.S., Aviles, F.X., Vendrell, J., and Ventura, S. (2006). Mutagenesis of the central hydrophobic cluster in Aβ42 Alzheimer’s peptide: Side-chain properties correlate with aggregation propensities. The FEBS Journal 273, 658–668. 10.1111/j.1742-4658.2005.05102.x.

16. Baumketner, A., Krone, M.G., and Shea, J.-E. (2008). Role of the familial Dutch mutation E22Q in the folding and aggregation of the 15–28 fragment of the Alzheimer amyloid-β protein. Proc. Natl. Acad. Sci. U.S.A. 105, 6027–6032. 10.1073/pnas.0708193105.

17. Faller, P. (2009). Copper and Zinc Binding to Amyloid-β: Coordination, Dynamics, Aggregation, Reactivity and Metal-Ion Transfer. ChemBioChem 10, 2837–2845. 10.1002/cbic.200900321.

18. Faller, P., and Hureau, C. (2009). Bioinorganic chemistry of copper and zinc ions coordinated to amyloid-β peptide. Dalton Trans., 1080–1094. 10.1039/B813398K.

19. Karr, J.W., Akintoye, H., Kaupp, L.J., and Szalai, V.A. (2005). N-Terminal Deletions Modify the Cu^2+^ Binding Site in Amyloid-β. Biochemistry 44, 5478–5487. 10.1021/bi047611e.

20. Miller, L.M., Wang, Q., Telivala, T.P., Smith, R.J., Lanzirotti, A., and Miklossy, J. (2006). Synchrotron-based infrared and X-ray imaging shows focalized accumulation of Cu and Zn co-localized with β-amyloid deposits in Alzheimer’s disease. Journal of Structural Biology 155, 30–37. 10.1016/j.jsb.2005.09.004.

21. Kawahara, M., Kato-Negishi, M., and Tanaka, K. (2020). Amyloids: Regulators of Metal Homeostasis in the Synapse. Molecules 25, 1441. 10.3390/molecules25061441.

22. Squitti, R., Faller, P., Hureau, C., Granzotto, A., White, A.R., and Kepp, K.P. (2021). Copper Imbalance in Alzheimer’s Disease and Its Link with the Amyloid Hypothesis: Towards a Combined Clinical, Chemical, and Genetic Etiology. JAD 83, 23–41. 10.3233/JAD-201556.

23. Jiang, D., Zhang, L., Grant, G.P.G., Dudzik, C.G., Chen, S., Patel, S., Hao, Y., Millhauser, G.L., and Zhou, F. (2013). The Elevated Copper Binding Strength of Amyloid-β Aggregates Allows the Sequestration of Copper from Albumin: A Pathway to Accumulation of Copper in Senile Plaques. Biochemistry 52, 547–556. 10.1021/bi301053h.

24. Karr, J.W., and Szalai, V.A. (2008). Cu(II) Binding to Monomeric, Oligomeric, and Fibrillar Forms of the Alzheimer’s Disease Amyloid-β Peptide. Biochemistry 47, 5006–5016. 10.1021/bi702423h.

25. Illodo, S., Al-Soufi, W., and Novo, M. (2024). Critical aggregation concentration and reversibility of amyloid-β (1–40) oligomers. Archives of Biochemistry and Biophysics 761, 110179. 10.1016/j.abb.2024.110179.

26. Novo, M., Freire, S., and Al-Soufi, W. (2018). Critical aggregation concentration for the formation of early Amyloid-β (1–42) oligomers. Sci Rep 8, 1783. 10.1038/s41598-018-19961-3.

27. Rudd, P. (2002). Glycosylation and prion protein. Current Opinion in Structural Biology 12, 578–586. 10.1016/S0959-440X(02)00377-9.

28. Salzano, G., Giachin, G., and Legname, G. (2019). Structural Consequences of Copper Binding to the Prion Protein. Cells 8, 770. 10.3390/cells8080770.

29. Fluharty, B.R., Biasini, E., Stravalaci, M., Sclip, A., Diomede, L., Balducci, C., La Vitola, P., Messa, M., Colombo, L., Forloni, G., et al. (2013). An N-terminal Fragment of the Prion Protein Binds to Amyloid-β Oligomers and Inhibits Their Neurotoxicity in Vivo. Journal of Biological Chemistry 288, 7857–7866. 10.1074/jbc.M112.423954.

30. Chen, S., Yadav, S.P., and Surewicz, W.K. (2010). Interaction between Human Prion Protein and Amyloid-β (Aβ) Oligomers. Journal of Biological Chemistry 285, 26377–26383. 10.1074/jbc.M110.145516.

31. Aronoff-Spencer, E., Burns, C.S., Avdievich, N.I., Gerfen, G.J., Peisach, J., Antholine, W.E., Ball, H.L., Cohen, F.E., Prusiner, S.B., and Millhauser, G.L. (2000). Identification of the Cu ^2+^ Binding Sites in the N-Terminal Domain of the Prion Protein by EPR and CD Spectroscopy. Biochemistry 39, 13760–13771. 10.1021/bi001472t.

32. Schilling, K.M., Tao, L., Wu, B., Kiblen, J.T.M., Ubilla-Rodriguez, N.C., Pushie, M.J., Britt, R.D., Roseman, G.P., Harris, D.A., and Millhauser, G.L. (2020). Both N-Terminal and C-Terminal Histidine Residues of the Prion Protein Are Essential for Copper Coordination and Neuroprotective Self-Regulation. Journal of Molecular Biology 432, 4408–4425. 10.1016/j.jmb.2020.05.020.

33. Evans, E.G.B., Pushie, M.J., Markham, K.A., Lee, H.-W., and Millhauser, G.L. (2016). Interaction between Prion Protein’s Copper-Bound Octarepeat Domain and a Charged C-Terminal Pocket Suggests a Mechanism for N-Terminal Regulation. Structure 24, 1057–1067. 10.1016/j.str.2016.04.017.

34. Wu, B., McDonald, A.J., Markham, K., Rich, C.B., McHugh, K.P., Tatzelt, J., Colby, D.W., Millhauser, G.L., and Harris, D.A. (2017). The N-terminus of the prion protein is a toxic effector regulated by the C-terminus. eLife 6, e23473. 10.7554/eLife.23473.

35. Jones, C.E., Klewpatinond, M., Abdelraheim, S.R., Brown, D.R., and Viles, J.H. (2005). Probing Copper2+ Binding to the Prion Protein Using Diamagnetic Nickel2+ and 1H NMR: The Unstructured N terminus Facilitates the Coordination of Six Copper2+ Ions at Physiological Concentrations. Journal of Molecular Biology 346, 1393–1407. 10.1016/j.jmb.2004.12.043.

36. Burns, C.S., Aronoff-Spencer, E., Legname, G., Prusiner, S.B., Antholine, W.E., Gerfen, G.J., Peisach, J., and Millhauser, G.L. (2003). Copper Coordination in the Full-Length, Recombinant Prion Protein. Biochemistry 42, 6794–6803. 10.1021/bi027138+.

37. Hooper, N.M., Taylor, D.R., and Watt, N.T. (2008). Mechanism of the metal-mediated endocytosis of the prion protein. Biochemical Society Transactions 36, 1272–1276. 10.1042/BST0361272.

38. Pauly, P.C., and Harris, D.A. (1998). Copper Stimulates Endocytosis of the Prion Protein. Journal of Biological Chemistry 273, 33107–33110. 10.1074/jbc.273.50.33107.

39. Atwood, C.S., Moir, R.D., Huang, X., Scarpa, R.C., Bacarra, N.M.E., Romano, D.M., Hartshorn, M.A., Tanzi, R.E., and Bush, A.I. (1998). Dramatic Aggregation of Alzheimer Aβ by Cu(II) Is Induced by Conditions Representing Physiological Acidosis. Journal of Biological Chemistry 273, 12817–12826. 10.1074/jbc.273.21.12817.

40. Walter, E.D., Chattopadhyay, M., and Millhauser, G.L. (2006). The Affinity of Copper Binding to the Prion Protein Octarepeat Domain: Evidence for Negative Cooperativity. Biochemistry 45, 13083–13092. 10.1021/bi060948r.

41. Millhauser, G.L., Singewald, K., Smart, A., and Pavlovici, F.A. (2026). Isotope-edited ESEEM: A new method for probing copper binding sites in neurodegenerative proteins. Journal of Biological Chemistry 302, 113310. 10.1016/j.jbc.2026.113310.

42. Dikanov, S., Felli, I., Viezzoli, M.-S., Spoyalov, A., and Hüttermann, J. (1994). X-band ESEEM spectroscopy of 15 N substituted native and inhibitor-bound superoxide dismutase: Hyperfine couplings with remote nitrogen of histidine ligands. FEBS Letters 345, 55–60. 10.1016/0014-5793(94)00406-4.

43. Lin, C.P., Bowman, M.K., and Norris, J.R. (1986). Analysis of electron spin echo modulation in randomly oriented solids: 15N modulation of radical cations of bacteriochlorophyll *a* and the primary donor of photosynthetic bacterium Rhodospirillumbrum. The Journal of Chemical Physics 85, 56–62. 10.1063/1.451582.

44. Quintanar, L., and Millhauser, G.L. (2022). EPR of copper centers in the prion protein. In Methods in Enzymology (Elsevier), pp. 297–314. 10.1016/bs.mie.2022.02.003.

45. Lee, H.-I., Doan, P.E., and Hoffman, B.M. (1999). General Analysis of 14N (I = 1) Electron Spin Echo Envelope Modulation. Journal of Magnetic Resonance 140, 91–107. 10.1006/jmre.1999.1803.

46. McCracken, J., Peisach, J., and Dooley, D.M. (1987). Cu(II) coordination chemistry of amine oxidases. Pulsed EPR studies of histidine imidazole, water, and exogenous ligand coordination. J. Am. Chem. Soc. 109, 4064–4072. 10.1021/ja00247a037.

47. Silva, K.I., Michael, B.C., Geib, S.J., and Saxena, S. (2014). ESEEM Analysis of Multi-Histidine Cu(II)-Coordination in Model Complexes, Peptides, and Amyloid-β. J. Phys. Chem. B 118, 8935–8944. 10.1021/jp500767n.

48. Clore, G.M., and Iwahara, J. (2009). Theory, Practice, and Applications of Paramagnetic Relaxation Enhancement for the Characterization of Transient Low-Population States of Biological Macromolecules and Their Complexes. Chem. Rev. 109, 4108–4139. 10.1021/cr900033p.

49. Donaldson, L.W., Skrynnikov, N.R., Choy, W.-Y., Muhandiram, D.R., Sarkar, B., Forman-Kay, J.D., and Kay, L.E. (2001). Structural Characterization of Proteins with an Attached ATCUN Motif by Paramagnetic Relaxation Enhancement NMR Spectroscopy. J. Am. Chem. Soc. 123, 9843–9847. 10.1021/ja011241p.

50. Kotler, S.A., Brender, J.R., Vivekanandan, S., Suzuki, Y., Yamamoto, K., Monette, M., Krishnamoorthy, J., Walsh, P., Cauble, M., Holl, M.M.B., et al. (2015). High-resolution NMR characterization of low abundance oligomers of amyloid-β without purification. Sci Rep 5, 11811. 10.1038/srep11811.

51. Danielsson, J., Pierattelli, R., Banci, L., and Gräslund, A. (2007). High-resolution NMR studies of the zinc-binding site of the Alzheimer’s amyloid β-peptide. The FEBS Journal 274, 46–59. 10.1111/j.1742-4658.2006.05563.x.

52. Pavlovici, F.A., Singewald, K., Kaplan, S., Chen, E., and Millhauser, G.L. (2025). Segmental Isotope Labeling of the Prion Protein: Identification of a Key Residue for Copper-Mediated Interdomain Structure. ACS Chem. Biol. 20, 1980–1989. 10.1021/acschembio.5c00336.

53. Gossert, A.D., Bonjour, S., Lysek, D.A., Fiorito, F., and Wüthrich, K. (2005). Prion protein NMR structures of elk and of mouse/elk hybrids. Proc. Natl. Acad. Sci. U.S.A. 102, 646–650. 10.1073/pnas.0409008102.

