## Supporting Information for "Bivalent, Copper-Mediated Amyloid-β Binding Disrupts the Prion Protein’s Neuroprotective Conformation"

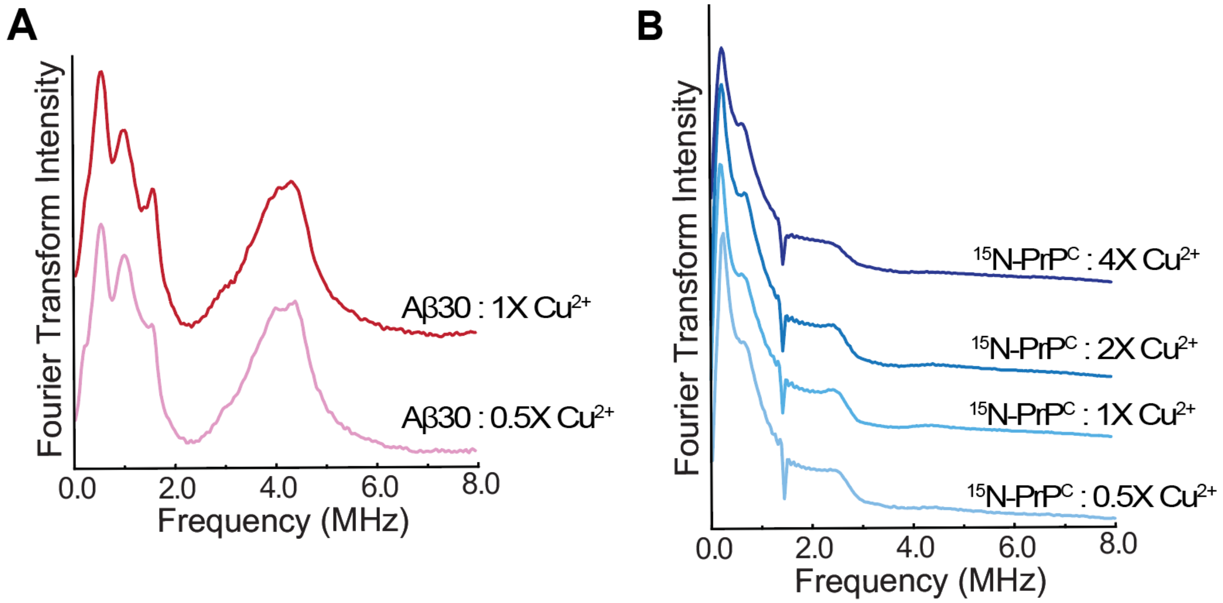


**Figure S1. ESEEM of Aβ30 and PrP$^{C}$ individually with increasing $Cu^{2+}$ concentration**. (**A**) The ESEEM frequency domain spectra of Aβ30 are shown for 0.5 and 1 eq. Concentrations beyond 1.0 eq. lead to free $Cu^{2+}$. (B) Stacked ESEEM frequency domain spectra of $^{15}$N-PrP$^{C}$ with 0.5, 1, 2 and 4 eq. of $Cu^{2+}$.

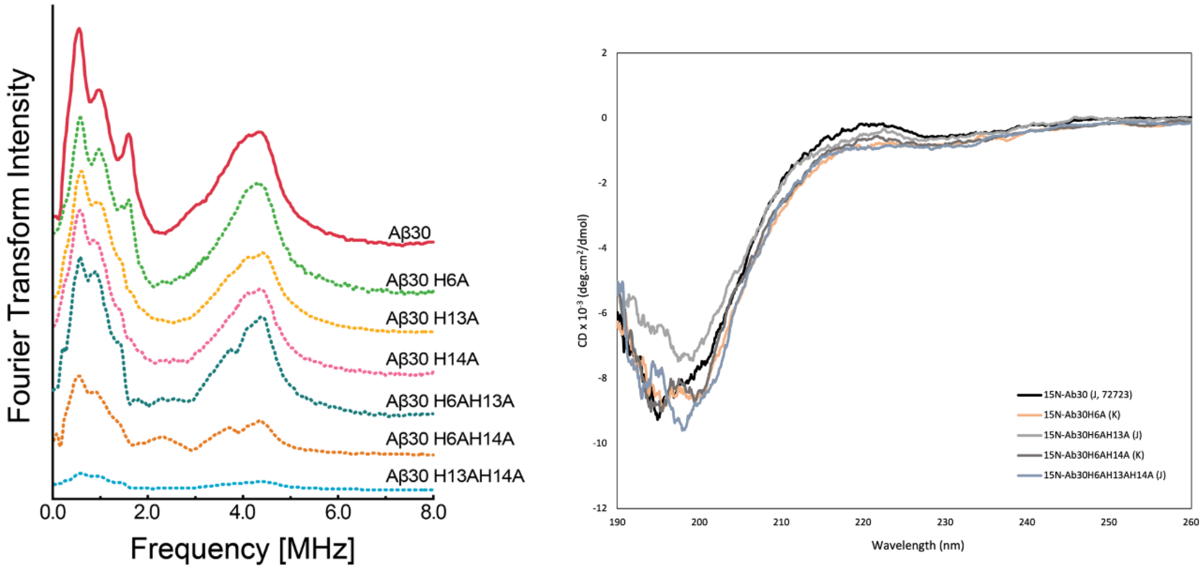


**Figure S2. ESEEM and CD of Aβ30 His mutants showing $Cu^{2+}$ coordination changes under NMR conditions.** The ESEEM frequency domain spectra (left) of Aβ30 mutants showing how any of the three His residues may coordinate to the $Cu^{2+}$ center although changes in the NQI region may reflect changes in local geometry. The CD spectra show random coil and no alpha helical nor beta sheet structure on 12.5 µM of $^{15}$N-Aβ30 (black), $^{15}$N-Aβ30 H6A (orange), $^{15}$N-Aβ30 H6A/H13A (grey), $^{15}$N-Aβ30H6A/H14A (blue), and $^{15}$N-Aβ30 H6A/H13A/H14A (light grey) at pH 6.6.

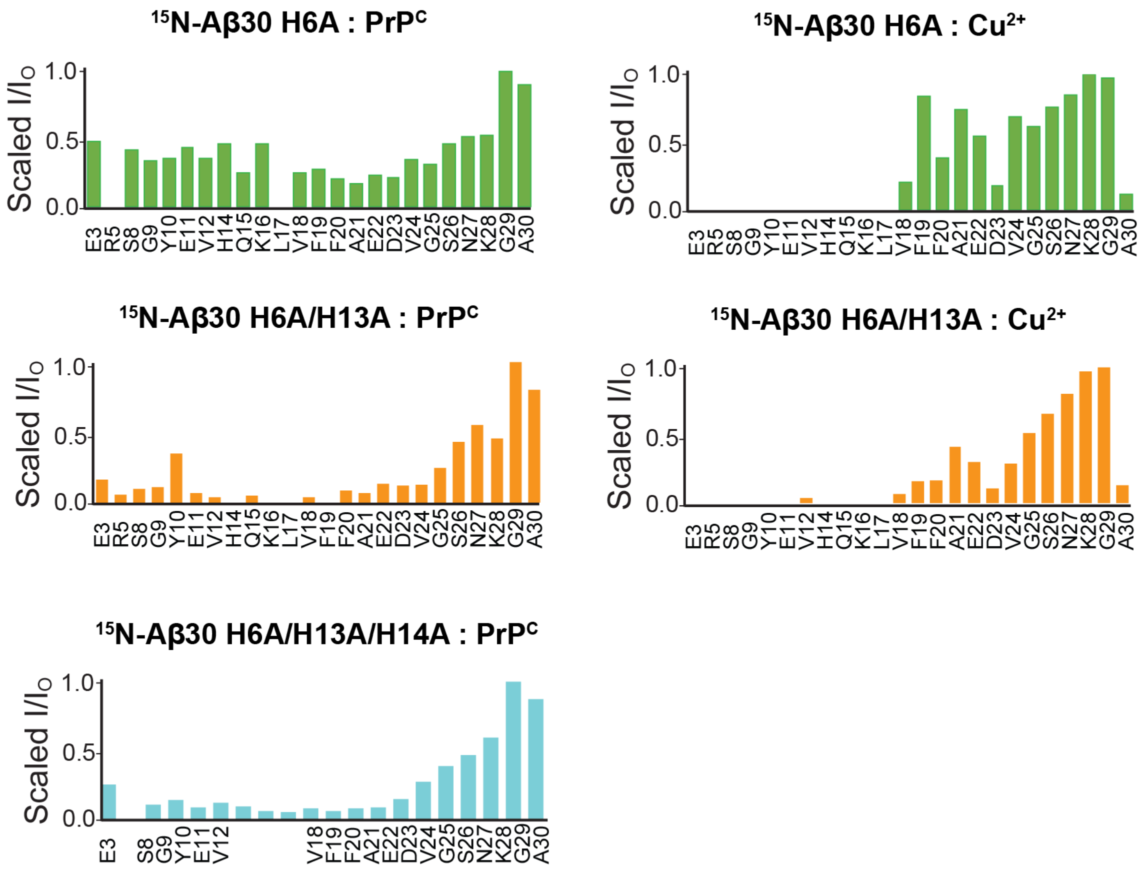


**Figure S3. The NMR intensity charts for Aβ30 His muts, with PrP$^{C}$ or Cu$^{2+}$.** Normalized intensity ratios from HSQC of $^{15}$N-Aβ30 H6A (green), $^{15}$N-Aβ30 H6A/H13A (orange), or $^{15}$N-Aβ30 ΔHis (cyan) with PrP$^{C}$ (left) or Cu$^{2+}$ (right). These show how Aβ30 His mutants change the interaction with PrP and Cu individually, outside of the ternary complex formation.

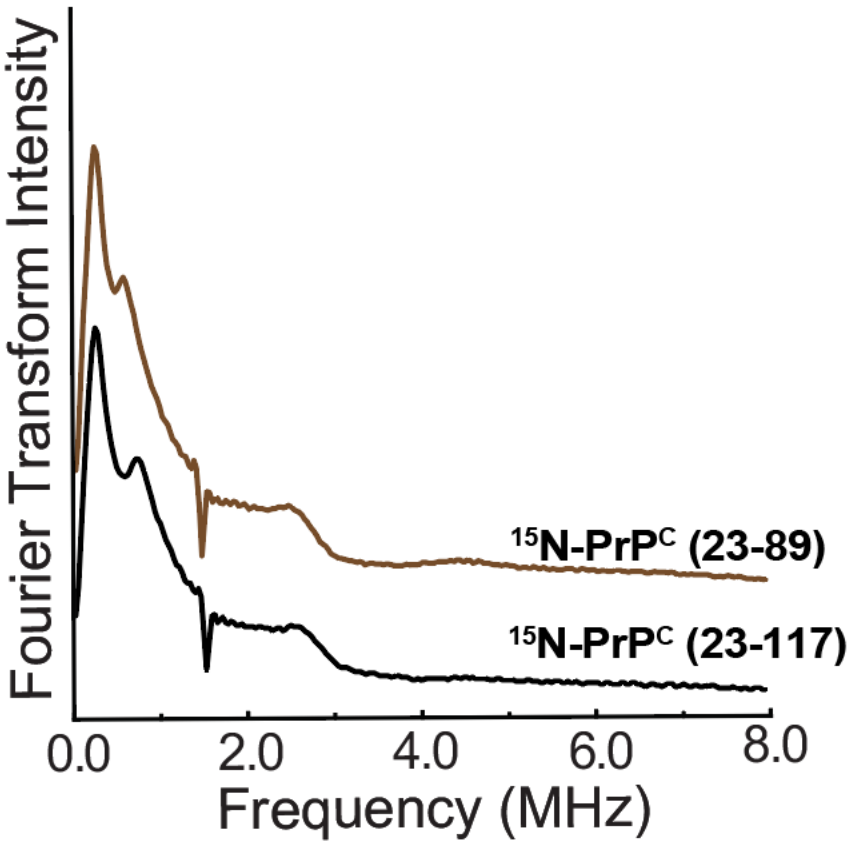


**Figure S4: ESEEM of $^{15}$N-PrP$^{C}$(23-89) and $^{15}$N-PrP$^{C}$(23-117) with $Cu^{2+}$.** Stacked ESEEM frequency domain spectra of $^{15}$N-PrP$^{C}$ (23-89) (top) and $^{15}$N-PrP$^{C}$ (23-117) (bottom) with 1 eq. $Cu^{2+}$. This shows how truncated PrP$^{C}$ variants coordinate $Cu^{2+}$.
